# Detecting Manuscripts Related to Computable Phenotypes Using a Transformer-based Language Model

**DOI:** 10.64898/2026.03.12.711165

**Authors:** Junghoon Chae, David Heise, Keith Connatser, Jacqueline Honerlaw, Monika Maripuri, Yuk-Lam Ho, Francesca Fontin, Vidisha Tanukonda, Kelly Cho

**Author notes:** **Corresponding Author** Junghoon Chae, Oak Ridge National Laboratory 1 Bethel Valley Road Oak Ridge, TN 37830.

## Abstract

**Objective:** The demand for a comprehensive phenomics library, which requires identifying computable phenotype definitions and associated metadata from an ever-expanding biomedical literature, presents a significant, labor-intensive, and unscalable challenge. To address this, we introduce a transformer-based language model specifically designed for identifying biomedical texts containing computable phenotypes and piloted its use in the Centralized Interactive Phenomics Resource (CIPHER) platform.

**Materials and Methods:** We fine-tuned a BioBERT model using a labeled dataset of 396 manuscripts. The model incorporates our novel sliding-window approach to effectively overcome token-length limitations, thereby enabling accurate classification of full-length manuscripts. For scalable deployment and continuous refinement, we developed a cohesive framework that integrates a web-based user interface, a control server, and a classification module.

**Results:** The staged approach for model development yielded a final model with 95% accuracy. The web-based user interface was deployed on the CIPHER platform and enables user feedback for model retraining.

**Discussion:** We developed a model and user interface which are currently in use by data curators to identify computable phenotype definitions from the literature.

**Conclusion:** Through this system, users can submit literature, assess classification results, and provide feedback directly influencing future model training, thereby offering an efficient and adaptive solution for accelerating phenotype-driven literature curation.

## BACKGROUND AND SIGNIFICANCE

Building a phenomics library – a comprehensive repository of computable phenotype definitions and associated metadata–relies heavily on the systematic extraction of relevant information from biomedical literature. One of the most critical and resource-intensive steps involves identifying content that contains sufficient information to support the development or recreation of computable phenotypes, which are typically found in published manuscripts. Given the growing biomedical literature, even experts struggle to locate relevant studies efficiently.

Natural Language Processing (NLP) has emerged as a promising solution for automating literature mining tasks, offering capabilities such as document classification, entity recognition, and relationship extraction. However, the effectiveness of many NLP models is often limited by their dependence on large, annotated datasets and significant computational resources. Widely used transformer-based models, such as Bidirectional Encoder Representations from Transformers (BERT) [1] and its variants, are typically constrained by a maximum input length of 512 tokens (where a token is roughly 3/4 of a word in English). This limitation is particularly problematic in biomedical research, where full-text articles often exceed 3,000 words, making it difficult to capture the contextual information necessary for accurate classification when only abstracts or partial texts are analyzed.

## OBJECTIVE

To address this, we developed a transformer-based model to identify full-length biomedical manuscripts relevant to computable phenotype development. This work also aimed to develop a framework and system implementation composed of a web-based user interface, a control server, and a classification module, which would not only integrate this transformer model, but also make it accessible for users to identify manuscripts and provide feedback for model retraining. The framework was implemented in the Centralized Interactive Phenomics Resource (CIPHER) platform, a public website which hosts a knowledgebase of phenotype definitions [2].

## MATERIALS AND METHODS

Our literature review framework is composed of four core components: a web-based user interface, a control server, storage module, and a classification module. The framework is illustrated in Figure 1. The web-based interface provides an accessible platform through which users can submit classification requests (e.g., by entering PubMed IDs) and view classification outputs such as relevance scores. It also allows users to provide feedback and enter metadata tags. This component emphasizes usability and transparency, allowing users to interpret outputs and contribute to refinement. The control server acts as the central coordinator within the framework. It is responsible for managing user interactions by receiving requests from the interface, forwarding them to the classification module, and delivering the resulting outputs back to the interface. The storage module manages most data such as feedback, tags, and comments from users. The classification module serves as the computational backbone, housing our model that performs manuscript classification. Upon receiving a request, it processes the input and returns the classification results. It also stores and manages user feedback to support periodic retraining, allowing the model to adapt to new data and evolving classification criteria. This design supports a dynamic learning process that enhances model accuracy over time.

**Figure 1.**
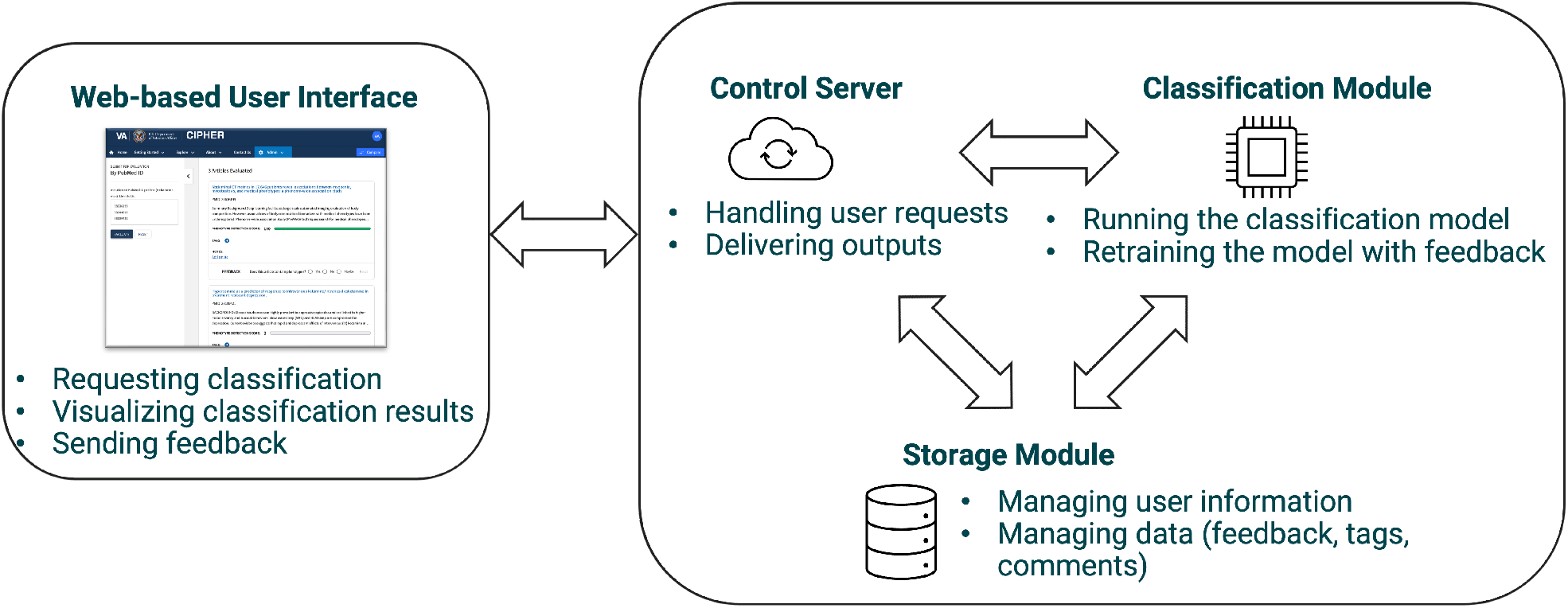
Comprehensive Systemic Framework for Computable Phenotype Literature Curation. This diagram illustrates the three modular components: the Web-based User Interface for submitting articles and viewing results, the Control Server for managing requests and responses, and the Classification Module for running the model and retraining based on user feedback. The arrows indicate the flow of information and interaction within the system.

### Data Preparation

Starting with about 176 biomedical research manuscripts, we progressively built a labeled dataset, eventually building a dataset of 396 manuscripts. Each document was manually reviewed and annotated by domain experts. Each document received a binary label indicating whether it contained sufficient information to support the recreation of a computable phenotype (“Yes”) or not (“No”). These annotations were made based on established criteria for phenotype reproducibility, including the presence of cohort definitions, inclusion/exclusion criteria, data sources, and algorithmic logic [3]. To ensure domain coverage and generalizability, the dataset spans a wide range of biomedical research areas, including cohort and population studies, electronic health record (EHR)-based phenotyping, clinical trials, and methodological papers.

### Classification Module

The classification module of the framework was developed through several iterative stages. We began with traditional machine learning algorithms suited to the initial data size and characteristics. As the dataset expanded and the documents required deeper contextual and semantic understanding, we transitioned to more advanced models. Each stage achieved notable improvements in classification performance, as summarized in Table 1. In this paper, we focus on the sliding-window model, which demonstrated the highest accuracy in the final stage.

**Table 1.**
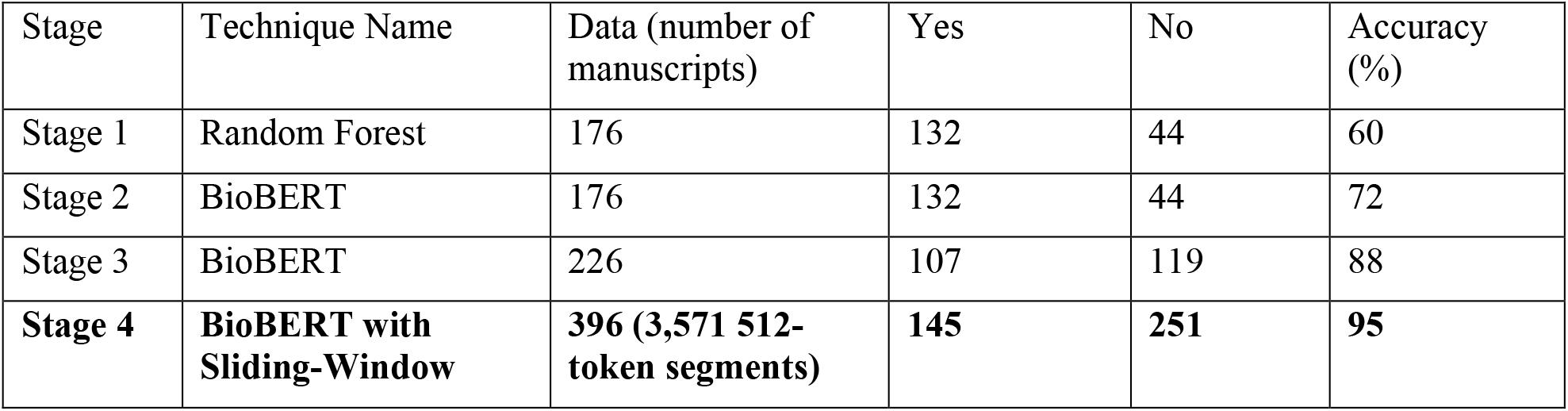
Progress of Performance Improvement.

| Stage | Technique Name | Data (number of manuscripts) | Yes | No | Accuracy (%) |
| --- | --- | --- | --- | --- | --- |
| Stage 1 | Random Forest | 176 | 132 | 44 | 60 |
| Stage 2 | BioBERT | 176 | 132 | 44 | 72 |
| Stage 3 | BioBERT | 226 | 107 | 119 | 88 |
| **Stage 4** | **BioBERT with Sliding-Window** | **396 (3,571 512-token segments)** | **145** | **251** | **95** |

#### Sliding-Window Method and Training

For the classification task, we employed BioBERT, a domain-specific adaptation of the BERT architecture [4]. BioBERT is pre-trained on large-scale biomedical corpora, including PubMed abstracts and full-text articles from PubMed Central (PMC). To address the token length limitation, we implemented a sliding-window segmentation approach. Each manuscript was divided into segments of 512 tokens. Given a tokenized document *D =* [*t*_1_, *t*_2_, …, *t*_*n*_],we divide it into non-overlapping segments of fixed length *L* (typically *L* = 512):

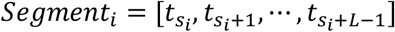

where the start index of each segment is:

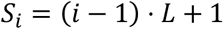

The number of segment *N* is:

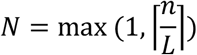

The binary label assigned to each original manuscript was propagated to all derived segments, creating a segment-level labeled dataset suitable for fine-tuning. This procedure expanded the original set of 396 documents into 3,571 labeled segments. Then, the dataset was randomly split into training and testing subsets using a 7:3 ratio. The model was fine-tuned on the segment-level training set using a binary cross-entropy loss function, optimizing the classification performance at the segment level.

#### Inference

At the inference step, each segment of a manuscript was independently classified by the fine-tuned model. To generate a manuscript-level prediction from these segment-level scores, we applied a weighted averaging aggregation strategy. Prediction probabilities for each segment are weighted based on segment length (the number of tokens) to account for varying information density. Let the model output a predicted probability *p*_*i*_ ∈ [0,1] for segment *i* and let *w*_*i*_ represents the weight of that segment, *w*_*i*_ = |*Segment*_*i*_|. The final document-level prediction score *P*_*D*_ is calculated as a weighted average:

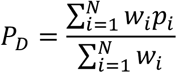

This approach ensures that longer, more content-rich segments exert greater influence on the final prediction while mitigating the impact of redundant or sparse text fragments.

### Web-Based User Interface

To support interactive evaluation of phenotype-relevant literature, we developed a web-based user interface. This interface allows users to submit articles for classification, review classification outputs, and provide feedback used to retrain the underlying model.

## RESULTS

Our model development progressed through several distinct stages, each contributing to significant improvements in classification accuracy and overall performance, as summarized in Table 1. Initially, our baseline was established using a Random Forest algorithm [5] (Stage 1). This traditional machine learning approach, trained on a biased dataset of over 176 manuscripts, yielded an accuracy of 60%. While providing a foundational benchmark, this result underscored the limitations of conventional methods for the complex task of biomedical literature classification.

The second stage introduced a transformer-based neural network model (Stage 2). By leveraging the power of transformers, even when trained on the same initial data, this model achieved a notable increase in accuracy to 72%. The receiver operating characteristic (ROC) curve for this stage in Figure 2 demonstrated improved performance in distinguishing between relevant and irrelevant manuscripts compared to the Random Forest baseline.

**Figure 2.**
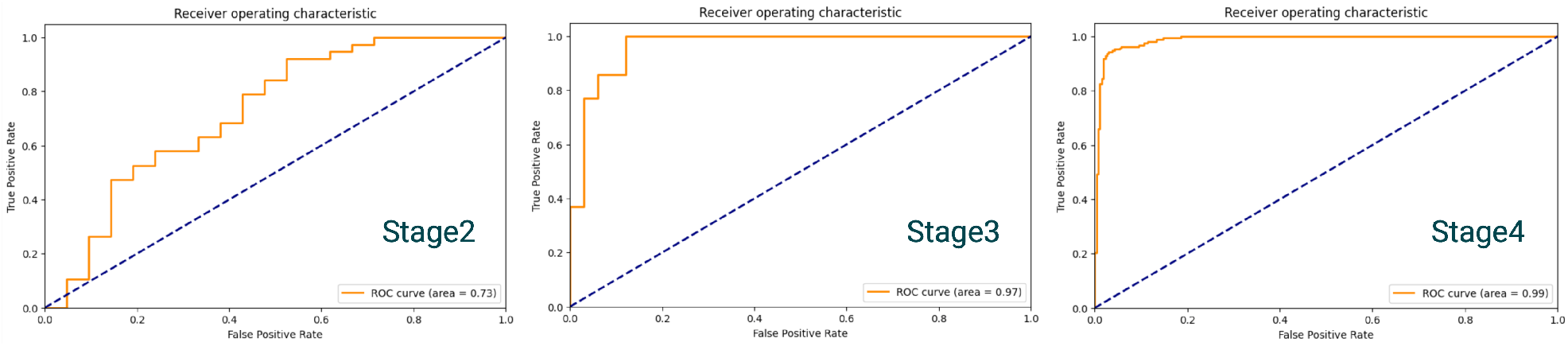
Receiver Operating Characteristic (ROC) Curves for Model Development Stages. The ROC curves illustrate the classification performance at different stages: Stage 2 (Transformer-based model with original data, Area Under Curve (AUC) = 0.72), Stage 3 (Transformer--based model with balanced data, AUC = 0.88), and Stage 4 (BioBERT with Sliding-Window approach, AUC = 0.99). The progression from left to right demonstrates the significant improvement in the true positive rate versus false positive rate with each model refinement.

We applied the same transformer-based technique to a new, balanced, dataset comprising 226 manuscripts (Stage 3). This strategic shift to a more representative and balanced dataset led to a jump in accuracy to 88%. The ROC curve for this iteration clearly shows a further upward and leftward shift, indicating enhanced sensitivity and specificity, and a more robust classification capability due to the improved data quality.

Finally, the most significant advancement was achieved with our sliding-window model (Stage 4), which incorporated a breakdown data approach. This method pushed the model’s performance to 95% accuracy. This improvement highlights the value of advanced architectures and the sliding-window approach in achieving high-accuracy classification for long biomedical texts.

The resulting model was integrated into the CIPHER platform. A screenshot of the interface is shown in Figure 3. The left-hand panel provides a submission area where users can enter PubMed IDs (PMIDs), with support for up to 5 entries at a time. Upon submission, the system retrieves and processes the corresponding manuscript. The system downloads full text articles if available through PMC and abstract only otherwise using PubMed APIs [6]. The main panel displays not only classification results, but also the titles and brief excerpts from the abstract, providing immediate context. A phenotype detection score is shown as a horizontal bar with a numerical value ranging from 0 to 100. This score reflects the model’s confidence in the presence of computable phenotype-relevant content.

**Figure 3.**
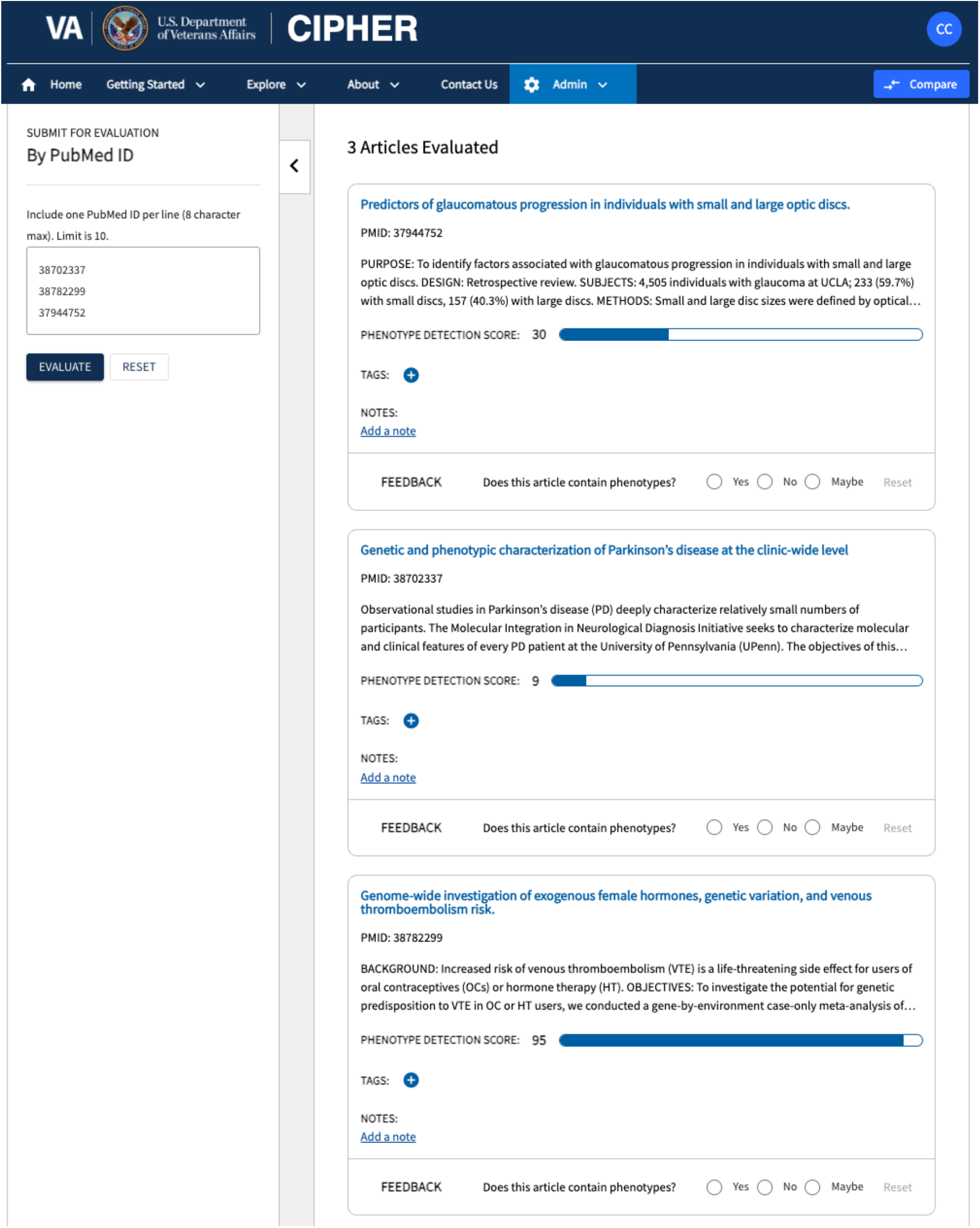
Screenshot of the Web-Based User Interface within the CIPHER Platform. This interface enables researchers to submit PubMed IDs for article classification, view predicted Phenotype Detection Scores, and provide feedback (Yes/No/Maybe) to continuously refine the underlying model. The left panel shows the submission area, and the main panel displays evaluation results for multiple articles.

The interface includes a feedback process where users are prompted to answer the question: “Does this article contain phenotypes?” with the options “Yes,” “No,” or “Maybe.” This label feedback is captured and stored in our backend database and is used for future model training. Users can optionally add *tags* and *notes* to each article for collaborative review. Currently, only “Yes” and “No” are used for model retraining.

### Use Case: CIPHER

This framework was developed for use within the CIPHER platform. Briefly, CIPHER is a knowledgebase of computable phenotype definitions developed by the United States Department of Veterans Affairs in partnership with the Department of Energy Oak Ridge National Laboratory. The knowledgebase and connected tools and resources are publicly accessible at https://phenomics.va.ornl.gov/. To grow the CIPHER knowledgebase, the program identifies computable phenotypes from published literature and manually adds phenotype metadata to the knowledgebase. The CIPHER team previously conducted literature reviews by manually identifying articles through targeted search criteria, which required significant time to review articles for relevance. With this web-based interface, the CIPHER team is now able to automate the initial review of abstracts for relevance using the Phenotype Detection Score, prioritizing our manual review efforts towards articles with a phenotype detection score of 50 or higher. As the model retraining continues, CIPHER plans to re-evaluate the threshold for manual review to further increase efficiency. Implementing use of this tool to filter out irrelevant manuscripts has enabled the CIPHER team to increase the number of publications reviewed and grow the phenotype metadata added to the library.

## DISCUSSION

We improved model performance in this specialized domain by addressing token-length limitations inherent in transformer models. However, deploying such a model alone does not resolve challenges related to scalability, user accessibility, and adaptability. Manual literature triage remains inefficient for large-scale or collaborative environments, and classification models must evolve continuously as new literature and user needs emerge. These limitations motivated the creation of a system that integrates the model with interactive feedback and retraining features.

The growing biomedical literature demands automated methods for information extraction and classification, particularly in medical informatics [7, 8]. Transformer-based models such as BioBERT [4], PubMedBERT [9], and ClinicalBERT [10] have shown strong performance in document classification but remain limited by a 512-token input cap caused by quadratic self-attention complexity. This restricts their direct application to long biomedical manuscripts. Existing strategies to overcome this issue either extend transformer capacity or segment documents. Models like Longformer [11] and Big Bird [12] use sparse attention mechanisms to process longer sequences efficiently, whereas methods such as Chunk-BERT [13], DocBERT [14], and ChunkBERT [15] divide texts into manageable parts.

The key difference in our work is applying a token-length–weighted aggregation strategy, ensuring that longer, information-dense segments have greater influence on the final document classification. Also, unlike BigBird or Chunk-BERT, our approach requires no model architecture modifications or post-processing pipelines, and our model is fully compatible with pre-trained biomedical models. This makes it well-suited for scalable, domain-specific applications such as phenotype-driven literature. Furthermore, our framework extends beyond just the classification model by incorporating a system that includes an interactive user interface and a control server that manages feedback for continuous model retraining, offering an adaptive and interactive solution for accelerating phenotype-driven literature curation. This framework for continuous model improvement and user accessibility is a crucial distinction from models that primarily focus on the classification algorithm itself.

Future work will focus on developing a Large Language Model (LLM) to automate the extraction of phenotypic information from scientific publications. Subsequently, we aim to establish a seamless pipeline to integrate this data into CIPHER. This initiative will streamline the current manual workflow, thereby enabling the team to prioritize high-level validation.

## CONCLUSION

This study developed and evaluated an automated approach to identify biomedical manuscripts relevant to computable phenotyping. This substantial performance gain highlights the method’s potential for scalable and highly efficient screening of biomedical literature, promising to considerably reduce the labor-intensive burden of manual literature curation and accelerate the identification of critical information for computable phenotype development.

## ACKNOWLEDGEMENTS

The contents do not represent the views of the U.S. Department of Veterans Affairs or the United States Government.

Notice: This manuscript has been authored by UT-Battelle, LLC, under contract DE-AC05-00OR22725 with the US Department of Energy (DOE). The US government retains and the publisher, by accepting the article for publication, acknowledges that the US government retains a nonexclusive, paid-up, irrevocable, worldwide license to publish or reproduce the published form of this manuscript, or allow others to do so, for US government purposes. DOE will provide public access to these results of federally sponsored research in accordance with the DOE Public Access Plan (https://www.energy.gov/doe-public-access-plan).

This research used resources at the Oak Ridge National Laboratory, which is supported by the Office of Science of the U.S. Department of Energy under Contract No. DE-AC05-00OR22725 and the Department of Veterans Affairs Office of Information Technology Inter-Agency Agreement with the Department of Energy under IAA No. VA118-16-M-1062.

